# Abstinence from Substance Use & The Value of Control

**DOI:** 10.1101/2025.05.02.651720

**Authors:** Elle M. Giovanni, Mimi Liljeholm

**Author notes:** **Funding:** National Science Foundation grant 1654187 (ML). National Institute on Drug Abuse training grant 1T32DA050558-01 (EG). **Declaration of interests:** Authors declare that they have no competing interests. **Data and materials availability:** All data and code are available on OSF. https://osf.io/yj7uk/?view_only=af1bc26fe6454bd6b85b2de05fb7ffde.

## Abstract

Clinical criteria for substance-use disorder collectively specify a loss of instrumental control over consumption, yet little empirical work has addressed how substance use shapes the utility of agency in motivated behavior. We combined a hierarchical gambling task with cross-sectional substance-use surveys and computational cognitive modeling, to assess the preference for controllable environments in adult humans with self-reported abstinence, across psychostimulant, opioid, alcohol, and sedative use. For psychostimulants, the duration of abstinence strongly modulated the preference for control, without affecting the ability to maximize monetary payoffs. In contrast, for alcohol, the recency of use was more predictive of a preference for environments with divergent outcome distributions, regardless of whether those outcomes were controllable. No effects of abstinence were seen for sedatives. The selective effects of psychostimulant use on the value of control implicate the dopaminergic system in agency coding.

## INTRODUCTION

Despite a longstanding debate regarding the legal and ethical implications of characterizing addiction as a voluntary choice (*1*), very little is known about how psychoactive substance use modulates the representation and utilization of agency. Healthy human adults (*2*–*6*), and indeed a range of non-human animals (*7*–*10*), have a well-demonstrated preference for free over forced choice. This preference may confer selective advantage: Since the subjective utilities of sensory stimuli (e.g., a particular food or piece of music) fluctuate continuously according to hedonic and homoeostatic dynamics, reward maximization requires an ability to flexibly produce distinct sensory states according to their current utilities (*11*–*14*). To better characterize the role of agency dysregulation in addiction, we assessed whether individual differences in the duration of self-reported abstinence from use predicted the subjective utility of instrumental control, across a range of psychoactive substances.

As an illustration of an environment’s controllability, consider the scenario depicted in Fig. 1A, where each of two gambling ‘rooms’ contains two options, and each option has a probability distribution over three outcome colors, as indicated by the pie charts. Assume that each color can be associated with any amount of monetary loss or gain, and that you must select a room to gamble in *before* the monetary color values are revealed, mimicking the volatility of subjective sensory utilities. In the left room, though afforded free choice, you would have very little control over which color, and thus which monetary outcome, was produced by your selections, since color outcome distributions are identical across options: Conversely, in the right room, though color distributions diverge across gambling options, you are forced to accept the selections of a computer algorithm that alternates between options across trials, again eliminating instrumental control.

**Figure 1.**
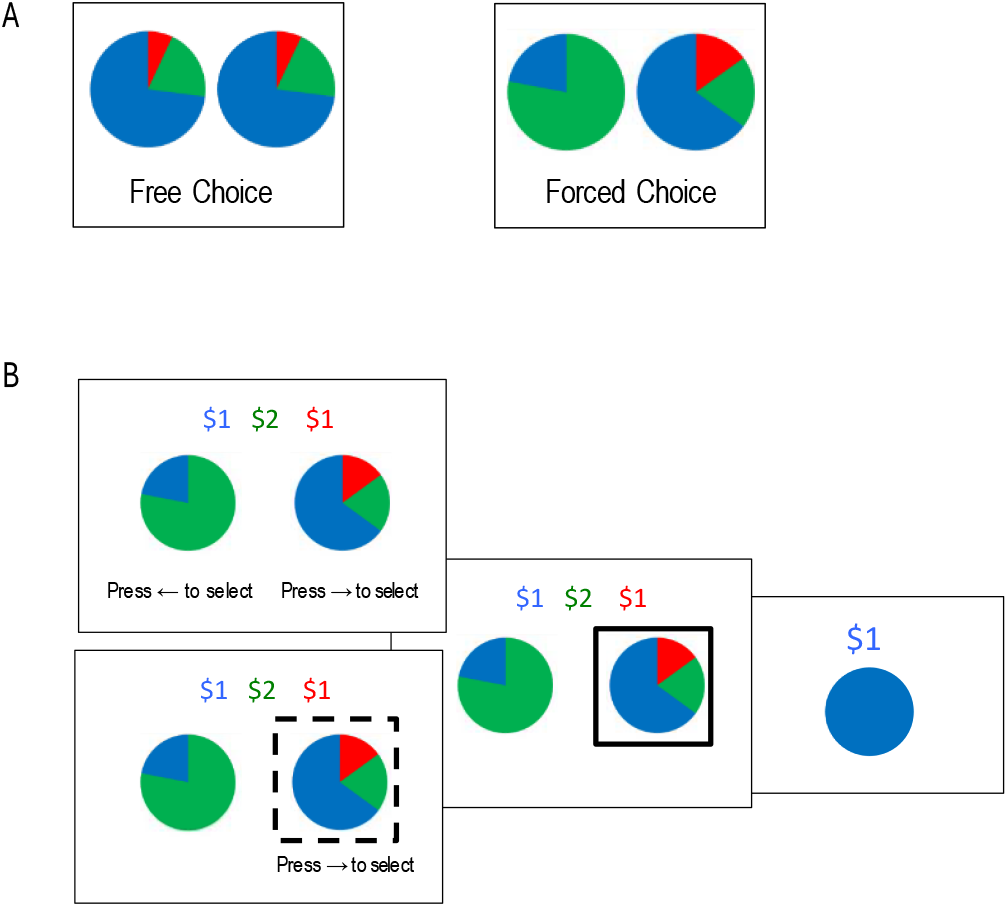
Task Illustration. A) Two gambling environments with low (Left) and high (Right) outcome divergence, with neither affording instrumental control. B) Choice, selection & feedback screens on a trial inside a gambling room, illustrating both free (top) and forced (bottom) choice screens. In Forced Choice rooms, dashed lines indicate the required selection.

When presented with choice scenarios such as that shown in Fig. 1A, neurotypical adults strongly prefer environments that are controllable: i.e., those in which freely chosen actions produce divergent outcome distributions (*11*–*14*). Here, we assess whether this preference for control is modulated by the recency of Psychostimulant, Opioid, Alcohol, and Sedative use.

## METHODS

### Participants

One hundred and thirty-three participants completed the study for monetary compensation (75 females; mean age = 38.4 ± 13.0 years). Recruitment aimed to obtain a sample size of at least 120, consistent with previous implementations of the task (*14*); it began on July 19, 2022, and ended on April 27, 2023. Each participant completed a single online session between July 22, 2022 and May 1, 2023. Participants were recruited using ResearchMatch – an NIH-funded online network that allows people with (and without) various medical conditions and life experiences to participate in IRB-approved studies at universities across the United States. We queried the database using addiction-related keywords (i.e., addiction, substance abuse) to identify potential participants.

### Task & Procedure

Our primary task measure was a commitment, at the start of each of several gambling rounds, to one of two ‘rooms’ in which to gamble for the duration of that round. The room-choice screens were as illustrated in Figure 1A, with pie charts representing the color outcome distributions of two alternative gambling options inside each room and with explicit labels indicating free vs forced choice, denoted in the study as ‘self-play’ and ‘auto-play’, respectively. Participants were instructed that they would be allowed to choose freely between options in ‘self-play’ rooms but that the computer would alternate between options across trials in ‘auto-play’ rooms. They were further told that the monetary values of color outcomes would change unpredictably across rounds (mimicking the fluctuating utilities of natural rewards) and only be revealed once a room had been selected. Their objective, thus, was to select, in each round, the gambling room that would allow them to maximize monetary payoffs in that round, given only information about the affordance of instrumental control, free choice, and sensory outcome divergence in each room.

Once a room had been selected, the monetary value of each color for that round was revealed, and four discrete gambling trials were performed in the room, with choice, selection, and feedback screens as illustrated in Figure 1B. Computer-selected gambling options were indicated by a dashed square outline, and participants had to press the corresponding key in order to progress through the trial. The ‘difference’ between outcome distributions associated with the gambling options inside a room was quantified as their information theoretic (Jensen-Shannon) divergence (*14*), with four divergences (0.00, 0.04, 0.15, & 0.20) yielding six unique differences in Outcome Divergence across gambling rooms. Specifically, each combination of divergences was combined with each self-vs. auto-play combination for a total of 24 unique room-choice scenarios, and these were repeated across two color schemes for 48 total room-choice trials and 192 total gambling trials. To ensure that stochastically generated monetary payoffs could not account for a controllability preference, reward distributions were constructed such that monetary payoffs were largely balanced and, if anything, biased against the hypothesized influence of control. To maintain incentive compatibility across rounds, participants were instructed that they would get to keep the monetary earnings, up to $15, from two gambling rounds, randomly drawn from all rounds at the end of the study.

### Computational Model

An environment’s controllability may be defined as the IT distance (*15*) between the outcome probability distributions associated with available actions. Let *P*_*1*_ and *P*_*2*_ be the respective color probability distributions of the two slot machines available in a given room, let *O* be the set of possible color outcomes, and *P(o)* the probability of a particular color outcome. The IT distance is:

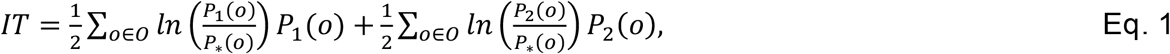

where *ln* is the natural logarithm, yielding nats, and

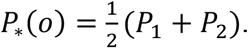

Note that, while an information theoretic divergence can be computed over distributions associated with any number and type of random variable(s), it only reflects instrumental control when computed over sensory outcome distributions associated with freely chosen actions (*14*)

An Expected Utility model specified a quantitative integration of the utility of Controllability with conventional monetary reward. First, the acquired value of a particular gambling room (i.e., a particular pair of pie charts) was incrementally updated across gambling rounds, such that

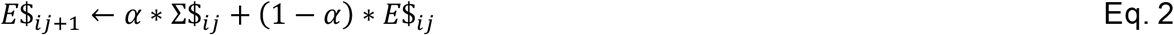

where *E*$_*ij*+1_ is the expected monetary pay-off in room *i* at the start of round *j+1* (i.e., at the start of the next round played in room *i*), Σ$_*ij*_ is the sum of monetary outcomes earned in room *i* at the end of round *j* (i.e., the end of the current round), *E*$_*ij*_ is the expected monetary payoff in room *i* on round *j* (recursively estimated based on the payoff, Σ$_*ij*−1_, and expectation, *E*$_*ij*−1_, in the previous round, *j-1*), and *α* is a free learning rate parameter.

The model further specified three free parameters: *γ*, indicating the subjective utility of Free Choice; *λ*, indicating the subjective utility of the Divergence, *D*, of color outcome distributions associated with the two options (i.e., pie charts) available in a room, and *β* indicating the subjective utility of Controllability (i.e., the combination of Free Choice and non-zero Outcome Divergence)

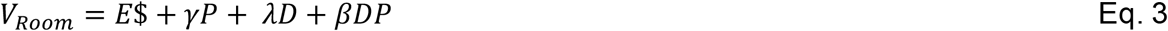

where *P* is an indicator set to 1 for Free Choice and 0 for Forced Choice.

Model-derived room values (*V*_*room*_) were transformed into room choice probabilities using a softmax rule with a noise parameter, *τ*;

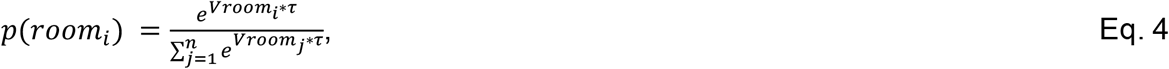

Free model parameters were fit to behavioral data by minimizing the negative log likelihood and computing the corrected Akaike Information Criterion (AIC). All computational variables were implemented using MATLAB (https://www.mathworks.com/).

### Statistical Analyses

Participants were grouped according to whether they had used a particular drug within the past 1 month (*Current* users) or abstained for 3 months or more (*Recovering* users) and a mixed Analyses of Variance (ANOVA) with Outcome Divergence and Free Choice as within-subjects variables, Current vs. Recovering users as a between-subject variable, and with usage of multiple drugs (henceforth poly-drug use), as covariates, was performed on the room-choice preferences. In addition, partial Spearman correlation coefficients, *ρ*, estimated the degree to which the duration of Abstinence from a particular drug predicted the amount of monetary gain obtained in controllable rooms and utility parameters derived from model fitting, again controlling for poly-drug use. All statistical analyses were performed in JASP (version 0.19.3) and all statistical outputs are listed in Supplemental Tables 1 and 2. Participation consisted of a single session during which all questionnaires and experimental tasks were completed, resulting in no missing data.

**Table 1:**
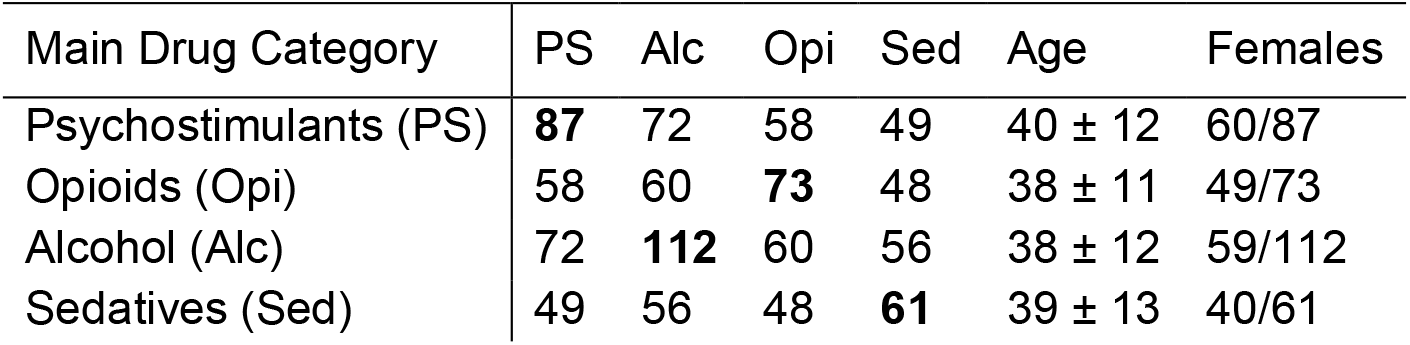
Sample demographics. Demographic information and number of individuals in each drug category that also had ever-use of drugs in one or more other category. The sample size for the Main Drug Category is in bold.

### Substance Use Assessment

Four categories of substances were screened: Psychostimulants (e.g., cocaine/crack, amphetamine), Opioids (e.g., heroin, fentanyl, hydrocodone), Sedatives (e.g., barbiturates, benzodiazepines), and Alcohol. The primary measure of interest was the duration of Abstinence from a particular drug of choice, indicated as currently using, one month, three months, six months, or more than one year of abstinence. In addition, scores were obtained for the duration of the last period of use (one month, three months, six months, 12 months, or longer than 12 months), and for use severity according to the *NIDA Quick Screen Version 1*, adapted to an online questionnaire format, for each substance.

## RESULTS

We identified and sent invitations to 4059 eligible volunteers by querying the RM database using addiction-related keywords (i.e., addiction, substance abuse. A total of 382 volunteers accepted the invitation to participate and were sent the study link, of which 133 completed all task elements in a single online session. Demographic and substance-use variables are listed in Table 1. All statistical outputs are listed in Supplemental Tables 1 and 2.

Of particular interest was how Abstinence from Psychostimulant, Opioid, Alcohol, and Sedative use related to preferences for gambling rooms as a function of Free Choice, Outcome Divergence, and Controllability. Mean choice preferences for gambling environments defined by Free Choice, Outcome Divergence, and Controllability, in Current (use in the last 1 month) and Recovering (no use for more than 3 months) users are plotted in Fig. 2, for each substance category. Strikingly, across substances, Current users do not display the characteristic preference for instrumental control clearly seen in Recovering users, as well as in previous studies using healthy volunteers (*14*). In particular, for Psychostimulants, Current users do not appear to discriminate between gambling environments based on either Free Choice or Outcome Divergence, while Recovering users clearly prefer Instrumental Control (i.e., *both* Free Choice & high Outcome Divergence).

**Figure 2.**
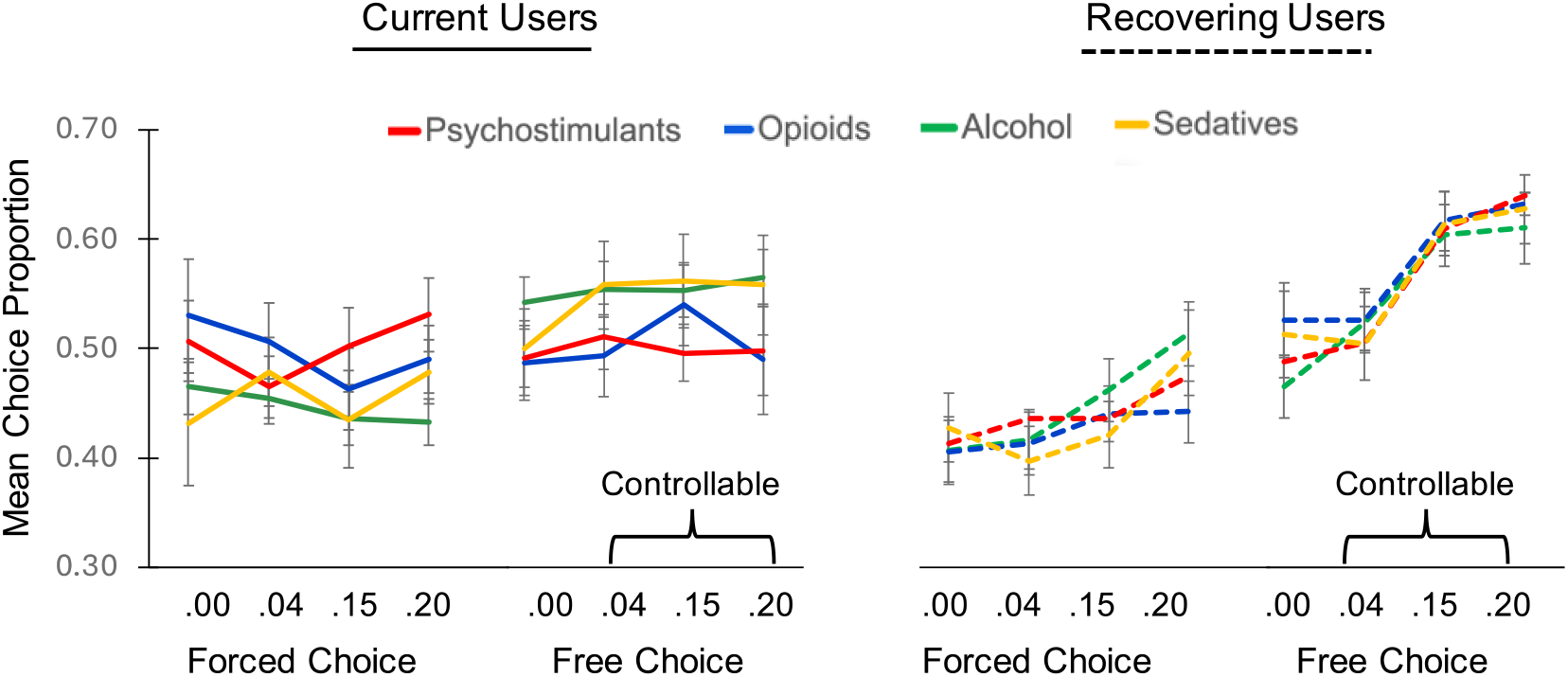
Mean choice preferences. Mean choice preferences across gambling environments as a function of Free Choice, Outcome Divergence, and Controllability in Current and Recovering users of each substance. Error bars = SEM.

Mixed ANOVAs were performed on choice preferences, separately for each drug category, with Free Choice and Outcome Divergence as within-subject factors, Abstinence as a between-subject factor, and with usage of each alternative drug category included as covariates. These analyses yielded significant Abstinence-by-Outcome Divergence-by-Free Choice interaction (*F*_3,246_ = 3.979, *p* = 0.009), as well as Abstinence-by-Free Choice interaction (*F*_*1,82*_ = 5.682, *p =* 0.019), for Psychostimulant use. For Opioid use, there was an Abstinence-by-Free Choice (*F*_1,68_ = 4.386, *p* = 0.04) interaction, and for Alcohol use an Abstinence-by-Outcome Divergence interaction (*F*_3,321_ = 3.724, *p* = 0.012). No significant effects involving Abstinence emerged for Sedative users (all p > 0.299).

For a more fine-grained analysis, we specified an expected utility model that generated choice preferences across gambling environments (see Methods). Briefly, the expected utility of a particular gambling room (i.e., a particular pair of pie charts) was a linear integration of its expected monetary payoff, acquired incrementally across rounds of gambling, and its affordance of Free Choice, Outcome Divergence, and Controllability, according to their respective subjective utilities.

As shown in Figure 3, partial Spearman correlation analyses, with ever-use of each alternative substance as partial correlates, revealed that longer Abstinence predicted a significantly greater model-derived subjective utility of Controllability exclusively for Psychostimulants (*ρ* = 0.226, *p* = 0.039, 95% CI [0.03, 0.43]). In Alcohol users, Abstinence predicted a greater subjective utility of Outcome Divergence (*ρ* = 0.224, *p* = 0.019, 95% CI [0.02, 0.42]). Critically, as shown in the right of Fig. 3, the ability to maximize monetary payoffs in controllable environments was only predicted by Abstinence in Sedative users (*ρ* = 0.270, *p* = 0.041, 95% CI [−0.01, 0.51]), suggesting that differences in the preference for Control across Current and Recovering Psychostimulant users was not due to impairments in basic reward processing.

**Figure 3.**
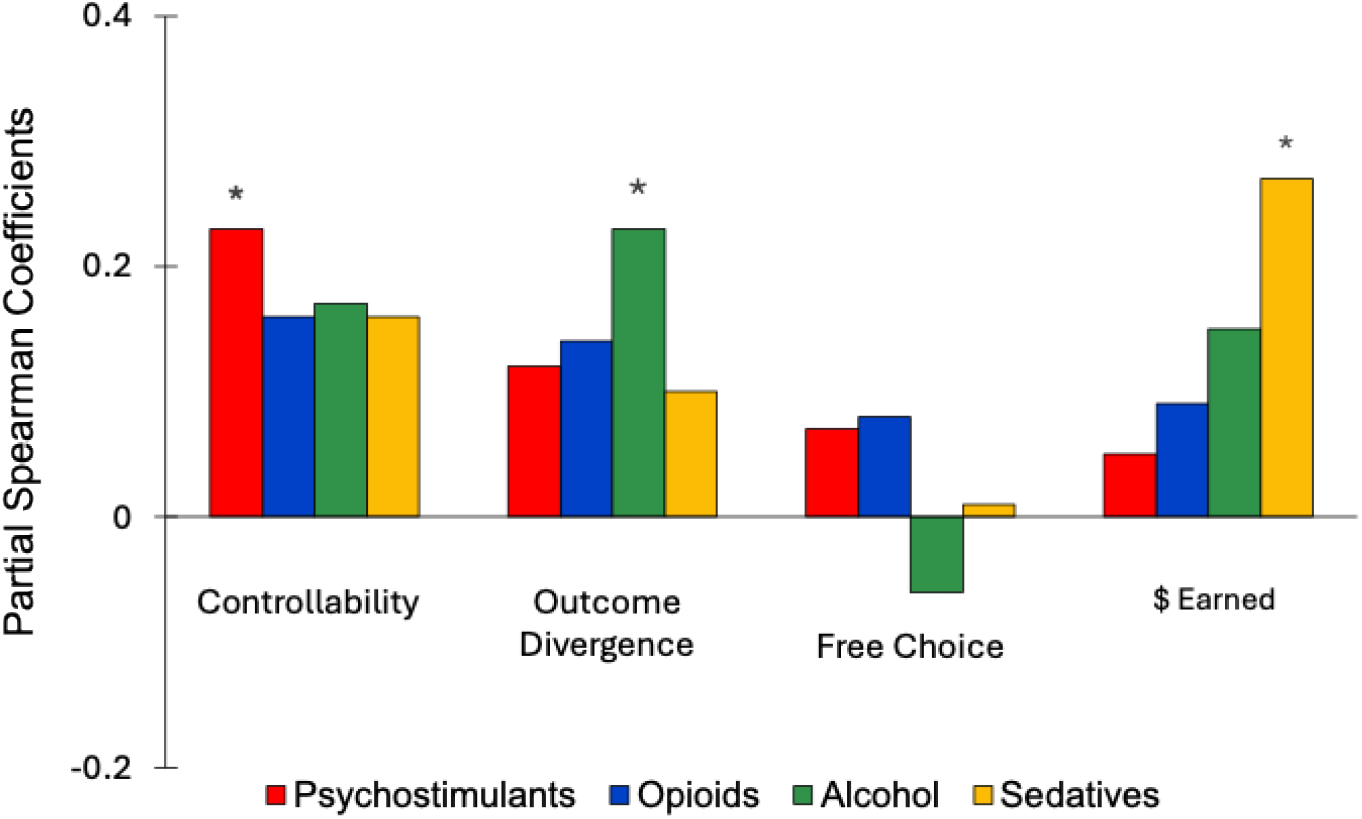
Partial Spearman correlation coefficients (*ρ*) estimating the relationship between Abstinence and variables of interest. Coefficients are plotted for correlations between Abstinence and computational cognitive model parameters, respectively estimating the subjective utilities of Controllability, Outcome Divergence and Free Choice, and between Abstinence and mean dollar amounts earned in controllable environments. \**p<*0.05

## DISCUSSION

We combined a hierarchical gambling task with cross-sectional substance use surveys and computational cognitive modeling to assess the preference for controllable environments in adult human participants with self-reported durations of abstinence from psychostimulant, opioid, alcohol, and sedative use. For psychostimulant use, abstinence strongly modulated the preference for controllable environments. In contrast, for alcohol, the recency of use was more predictive of a preference for environments with divergent outcome distributions across available options, regardless of whether those outcomes were controllable. For opioids, results were equivocal, with a significant effect of abstinence on the free-choice preference emerging in exploratory analyses, but not in model-based parameter estimations. Finally, for sedative use, the only significant effect of abstinence was on the ability to maximize monetary reward; critically, this measure of reward maximization was unaffected by abstinence from psychostimulant, opioid, and alcohol use, dissociating control, and its subcomponents, from basic reward processes.

Since the subjective utilities of sensory states fluctuate constantly according to hedonic and homeostatic principles, reward maximization depends critically on the ability to flexibly produce or prevent states according to their current utilities. Consistent with this notion, neurotypical human decision-makers prefer, all else being equal, to act in environments with greater controllability, often formalized as the information theoretic distance between sensory distributions associated with freely chosen actions (*11, 14*). This preference, observed here in recovering, but not current psychostimulant users, has been shown to scale with activity in the ventromedial prefrontal cortex (*16*), an area that is robustly implicated in domain general decision-value computations (*17*–*19*), and to decrease with *positive schizotypy:* a personality trait characterized by unusual thoughts and experiences (*14, 20*).

The reduced preference for controllability for both positive schizotypy and psychostimulant use might reflect a shared dopaminergic (DA) basis. Several studies have identified aberrant DA transmission in positive schizotypy (see (*21*) for review), and DA imbalances are well documented in its clinical counterpart, schizophrenia (*22, 23*). Unlike Opioids and Alcohol, Psychostimulants like amphetamine and cocaine act on a range of presynaptic mechanisms to increase extracellular DA, including blocking the transport of DA back into the presynaptic cell, and mobilizing vesicular DA release (*24, 25*). Notably, Schizotypy, Psychostimulants, and baseline DA levels, have all been associated with aberrant self vs external control attributions (*26*–*29*). Together, these results support the notion of a dopaminergic agency code (*30, 31*).

This work has identified several core distinctions with respect to the influence of substance-use on the preference for instrumental control, including that between different illicit substances, and corollary neuromodulatory systems, between dissociable sub-components of instrumental control, such as free choice and outcome divergence, and between the utility of control and basic monetary reward maximization. The study also suffers from several limitations, including the lack of clear separation between substances, which we have addressed by including poly-use in our statistical models, the assessment of abstinence duration as broad categories rather than months, reducing the sensitivity of the measure, and the lack of a comprehensive independent assessment of addiction severity. Future studies will need to address these factors in order to further characterize the role of instrumental control, and agency more broadly, in substance use disorder.

