## Supplemental Tables 1-2 for "Abstinence from Substance Use & The Value of Control"

### SUPPLEMENTAL MATERIALS

#### Tables

*Supplemental Table 1.* Statistical output from Mixed ANOVAs, one per substance, assessing the influence of Abstinence, Outcome Divergence (OD), Free Choice (FC), & Polysubstance Use for each alternative drug category, on Gambling Environment Preferences.

| Main Drug Category | Effect | df | F | *p* |
|---|---|---|---|---|
| Psychostimulants | Free Choice (FC) | 1,82 | 1.13 | 0.29 |
| | FC x Abstinence | 1,82 | 5.68 | 0.02 |
| | FC x Opioid Use | 1,82 | 0.07 | 0.79 |
| | FC x Alcohol Use | 1,82 | 3.48 | 0.07 |
| | FC x Sedative Use | 1,82 | 0.11 | 0.74 |
| | Outcome Divergence (OD) | 3,246 | 0.16 | 0.92 |
| | OD x Abstinence | 3,246 | 1.60 | 0.20 |
| | OD x Opioid Use | 3,246 | 2.05 | 0.11 |
| | OD x Alcohol Use | 3,246 | 0.57 | 0.64 |
| | OD x Sedative Use | 3,246 | 1.59 | 0.20 |
| | FC x OD | 3,246 | 1.02 | 0.39 |
| | FC x OD x Abstinence | 3,246 | 3.98 | 0.01 |
| | FC x OD x Opioid Use | 3,246 | 4.35 | <0.01 |
| | FC x OD x Alcohol Use | 3,246 | 0.55 | 0.65 |
| | FC x OD x Sedative Use | 3,246 | 1.44 | 0.23 |
| Alcohol | FC | 1,107 | 5.64 | 0.02 |
| | FC x Abstinence | 1,107 | <0.01 | 0.98 |
| | FC x Psychostimulant Use | 1,107 | 0.85 | 0.36 |
| | FC x Opioid Use | 1,107 | 0.59 | 0.45 |
| | FC x Sedative Use | 1,107 | 0.01 | 0.91 |
| | OD | 3,321 | 0.81 | 0.49 |
| | OD x Abstinence | 3,321 | 3.72 | 0.01 |
| | OD x Psychostimulant Use | 3,321 | 1.74 | 0.16 |
| | OD x Opioid Use | 3,321 | 0.63 | 0.60 |
| | OD x Sedative Use | 3,321 | 0.15 | 0.93 |
| | FC x OD | 3,321 | 0.78 | 0.51 |
| | FC x OD x Abstinence | 3,321 | 0.43 | 0.74 |
| | FC x OD x Psychostimulant Use | 3,321 | 0.05 | 0.99 |
| | FC x OD x Opioid Use | 3,321 | 3.40 | 0.02 |
| | FC x OD x Sedative Use | 3,321 | 0.79 | 0.50 |
| Opioids | FC | 1,68 | 1.24 | 0.27 |
| | FC x Abstinence | 1,68 | 4.39 | 0.04 |
| | FC x Psychostimulant Use | 1,68 | 3.08 | 0.08 |
| | FC x Alcohol Use | 1,68 | 0.35 | 0.56 |
| | FC x Sedative Use | 1,68 | 1.23 | 0.27 |
| | OD | 3,204 | 0.11 | 0.95 |
| | OD x Abstinence | 3,204 | 0.84 | 0.47 |
| | OD x Psychostimulant Use | 3,204 | 0.08 | 0.97 |
| | OD x Alcohol Use | 3,204 | 0.05 | 0.98 |
| | OD x Sedative Use | 3,204 | 1.62 | 0.19 |

| | | | | |
|---|---|---|---|---|
| | FC x OD | 3,204 | 0.42 | 0.74 |
|  | FC x OD x Abstinence | 3,204 | 1.01 | 0.39 |
| | FC x OD x Psychostimulant Use | 3,204 | 1.70 | 0.17 |
| | FC x OD x Alcohol Use | 3,204 | 0.82 | 0.48 |
| | FC x OD x Sedative Use | 3,204 | 0.71 | 0.55 |
| Sedatives | FC | 1,56 | 2.06 | 0.16 |
|  | FC x Abstinence | 1,56 | 0.92 | 0.34 |
| | FC x Psychostimulant Use | 1,56 | 2.26 | 0.14 |
| | FC x Alcohol Use | 1,56 | 0.89 | 0.35 |
| | FC x Opioid Use | 1,56 | 0.25 | 0.62 |
| | OD | 3,168 | 1.77 | 0.16 |
|  | OD x Abstinence | 3,168 | 1.24 | 0.30 |
| | OD x Psychostimulant Use | 3,168 | 1.19 | 0.32 |
| | OD x Alcohol Use | 3,168 | 0.30 | 0.83 |
| | OD x Opioid Use | 3,168 | 2.52 | 0.06 |
| | FC x OD | 3,168 | 0.61 | 0.61 |
|  | FC x OD x Abstinence | 3,168 | 0.11 | 0.95 |
| | FC x OD x Psychostimulant Use | 3,168 | 0.16 | 0.92 |
| | FC x OD x Alcohol Use | 3,168 | 2.68 | 0.05 |
| | FC x OD x Opioid Use | 3,168 | 0.13 | 0.94 |

*Supplemental Table 2.* Statistical output from Partial Spearman Correlations, assessing the relationship between Abstinence and model parameters estimating the subjective utility of Free Choice, Outcome Divergence, and Controllability, as well as between Abstinence and mean monetary earnings in controllable rooms, controlling for ever-use of each alternative substance.

| | Substance | $\rho$ | $p$ | 95% CI |
|---|---|---|---|---|
| Free Choice | Psychostimulants | 0.07 | 0.54 | [−0.14, 0.30] |
| | Alcohol | -0.06 | 0.53 | [−0.24, 0.14] |
| | Opioids | 0.08 | 0.54 | [−0.16, 0.31] |
| | Sedatives | 0.01 | 0.93 | [−0.27, 0.27] |
| Outcome Divergence | Psychostimulants | 0.12 | 0.29 | [−0.08, 0.32] |
| | Alcohol | 0.23 | 0.02 | [0.02, 0.42] |
| | Opioids | 0.14 | 0.26 | [−0.08, 0.35] |
| | Sedatives | 0.10 | 0.44 | [−0.15, 0.35] |
| Controllability | Psychostimulants | 0.23 | 0.04 | [0.03, 0.43] |
| | Alcohol | 0.17 | 0.08 | [−0.01, 0.35] |
| | Opioids | 0.16 | 0.18 | [−0.07, 0.38] |
| | Sedatives | 0.16 | 0.22 | [−0.13, 0.42] |
| $ Earned | Psychostimulants | 0.05 | 0.63 | [−0.16, 0.26] |
| | Alcohol | 0.15 | 0.12 | [−0.07, 0.33] |
| | Opioids | 0.09 | 0.44 | [−0.11, 0.32] |
| | Sedatives | 0.27 | 0.04 | [−0.01, 0.51] |
